# Novel Real-Time Library Search driven data acquisition strategy for identification and characterization of metabolites

**DOI:** 10.1101/2021.10.07.463419

**Authors:** Brandon Bills, William D. Barshop, Seema Sharma, Jesse Canterbury, Aaron M. Robitaille, Michael W. Senko, Vlad Zabrouskov

## Abstract

Identification and structural characterization of novel metabolites in drug discovery or metabolomics experiments is one of the most challenging tasks. Multi-level fragmentation (MS^n^) based approaches combined with various dissociation modes are frequently utilized for facilitating structure assignment of the unknown compounds. As each of the MS precursors undergoes MS^n^, the instrument cycle time can limit the total number of precursors analyzed in a single run for complex samples. This necessitates splitting data acquisition into several LC/MS analyses where the results obtained in one acquisition inform the experimental design for the successive experiment.

Here we present a new LC/MS data acquisition strategy, termed Met-IQ, where the decision to perform an MS^n^ acquisition is automatically made in real time based on the similarity between an acquired experimental MS^2^ spectrum and a spectrum in a reference spectral library. Each MS^2^ spectrum is searched in real time against the spectra for the known compounds of interest. If a similarity to a spectrum in the library is found, the instrument performs a decision-dependent event, such as an MS^3^ scan. Compared to an intensity-based, data-dependent MS^n^ experiment, only a selective number of MS^3^ are triggered using Met-IQ, increasing the overall MS^2^ instrument sampling rate. We applied this strategy to an Amprenavir sample incubated with human liver microsomes. The number of MS^2^ scan events increased nearly 3.5-fold compared to the standard data dependent experiment where MS^3^ was triggered for each precursor ion, resulting in identification and structural characterization of 55% more unique metabolites. Furthermore, the MS^3^ precursor fragments were selected intelligently, focusing on higher mass fragments of sufficient intensity to maximize acquisition of MS^3^ data relevant for structure assignment. The described Met-IQ strategy is not limited to metabolism experiments, and can be applied to analytical samples where the detection of unknown compounds structurally related to a known compound(s) is sought.

## Introduction

In applications such as drug development or environmental safety, metabolites need to be fully characterized to investigate any toxic effects. Mass spectrometry paired with liquid chromatography is often the analytical method of choice; however, characterizing these metabolites can be challenging^1^. Targeted analysis can miss some metabolites because it requires a comprehensive list of potential metabolites and it can be difficult to predict exactly how a compound will degrade or be metabolized in a complex biological system. Untargeted analysis can acquire more comprehensive data by interrogating as many compounds as possible in a single run, but this leads to cumbersome analysis of large data sets and low concentration metabolites can be missed when high concentration background compounds from the matrix take up all the cycle time.

Modern mass spectrometers (MS) have built-in data acquisition strategies to intelligently guide the instrument. Filters can be implemented to only consider compounds of sufficient intensity or to target compounds exhibiting a relevant neutral loss. Workflows such as the Thermo Scientific™ AcquireX data acquisition workflow can direct the instrument to automatically analyze a matrix blank and add those compounds to an exclusion list to allow the instrument to focus on analyzing likely compounds of interest^2^. These techniques significantly improve the chances of detecting every metabolite but can require multiple analytical runs. In addition, a static instrument method may not capture enough information on every compound of interest for full structural characterization.

Real-time decision making allows the instrument to change the scan behavior based on the data being acquired during the analytical run. The software takes acquired data, compares it against a database and uses that information to choose a follow up scan during that same analysis. In the field of proteomics, specifically peptide quantitation using isobaric tags, Real-Time Search is used to avoid wasting instrument time by comparing peptide MS^2^ spectra to an in-silico peptide database to trigger MS^3^ scans only on the confidently identified peptides^3,4^. Here we introduce a small molecule focused, real-time spectral library search (RTLS)-based data acquisition strategy, on the Thermo Scientific™ Orbitrap IQ-X™ Tribrid™ MS termed Met-IQ. For small molecule experiments, the focus is on directing the instrument to acquire additional data on poorly identified compounds. Experimental fragmentation spectra are compared to a spectral library, such as an offline version of Thermo Scientific mzCloud™ library or a curated spectra from Thermo Scientific™ mzVault™ library^5^, and based on the result, the instrument can trigger additional scan behavior, such as MS^n^ or alternate fragmentation techniques, to get additional structural information about the compound. The new Met-IQ data acquisition strategy uses RTLS to compare incoming MS^2^ spectra against the spectra of the unmetabolized compound, regardless of whether the experimental precursor *m/z* matches the library precursor *m/z*, to check for spectral similarity. If spectral similarity is established between the spectrum for an unknown compound and the library spectrum of unmetabolized compound, meaning there are sufficient matching MS^2^ fragments to meet the criteria set in the method editor, then the instrument is automatically directed to perform MS^3^ scans on such putative metabolites. A sample of the drug Amprenavir metabolized by human liver microsome was analyzed with and without the Met-IQ data acquisition strategy to determine whether the RTLS-guided method improved the detection of metabolites in a single run over an unguided analytical approach.

## Experimental

### Creating the Library

Analyses were carried out on a sample of metabolized Amprenavir with and without Real-Time Library Search then compared for improvements in metabolite profiling. In order to use the Met-IQ data acquisition strategy, an MS^2^ library had to be generated for the unmetabolized form of the drug. A 250 ng/mL solution of Amprenavir in 50% methanol 50% water with 0.1% formic acid was infused on the Orbitrap IQ-X Tribrid mass spectrometer. MS^2^ were collected in positive mode using HCD at collision energies 10, 20, 30, 40, 50, 60, 70, and 80 as well as CID at collision energies 15, 30, and 45. Spectra were compiled into a database using mzVault software.

### Configuring Real-Time Library Search

Within the method editor, the Met-IQ template was selected then the RTLS node was modified for the analysis of Amprenavir. The spectral library created in mzVault was selected through the “Browse” feature. The value for the cosine score threshold, the score representing matched peaks and peak intensities between the experimental and library MS^2^ spectra, was increased to 15 to reduce the number of false positive triggers.

### Analysis

A sample of 5 μM Amprenavir was digested by incubation with human liver microsome with NADPH and GSH. UHPLC-MS analysis was carried out on a Thermo Scientific™ Vanquish™ Horizon UHPLC system coupled to an Orbitrap IQ-X Tribrid mass spectrometer. UHPLC was carried out in reverse phase mode using a gradient elution with an aqueous phase of water with 0.1% formic acid and an organic phase of acetonitrile with 0.1% formic acid. Analytical runs were 18 minutes long with a total of 5 μL of sample injected on the column. Mass spectra were collected in positive ion mode using data dependent MS^2^ (ddMS^2^) with stepped HCD activation at normalized collision energies of 20, 40, and 60%. In addition, ddMS^3^ was collected using two different methods, a control using only data dependent acquisition (DDA) and the same method but using Real-Time Library Search to guide triggering of ddMS^3^.

### Data Processing

Data was processed using Thermo Scientific™ FreeStyle™ software. Known metabolites of Amprenavir were manually checked within the list of MS^2^ and MS^3^ spectra. A compound was considered a potential metabolite if the precursor *m/z* matched a known metabolite within 3 ppm and there was at least one matched fragment in common between the experimental spectrum and the spectral library. In addition, the total number of MS^2^ and MS^3^ spectra was tabulated. Compounds were further investigated using Thermo Scientific™ Compound Discoverer™ software.

## Results and Discussion

### Real-Time Library Search Architecture

The Real-Time Library Search for small molecules on the Orbitrap IQ-X Tribrid MS utilizes a shared underlying architectural framework with the peptide Real-Time Search feature as implemented on the Orbitrap Eclipse Tribrid MS instrument^3^. At the start of each method execution, as shown in Figure 1, the user specified RTLS filter parameters from the method are transferred to the Real-Time Search service on the Data System PC. First, the Real-Time Search service will load the provided mzVault (.db) library into an in-memory *m/*z indexed structure, while filtering the library contents based on the method design. Library spectra which do not match the RTLS method MS^2^ acquisition settings for polarity, activation mode, and analyzer type are omitted from inclusion in the scorable candidate index. Next, the Real-Time Search service signals its ready status to the instrument firmware and awaits scans from method acquisition. MS^2^ spectra generated on the instrument from the relevant scan node to which the RTLS filter node is attached are first serialized into FlatBuffers prior to being sent from the instrument firmware to the Real-Time Search service.

**Figure 1:**
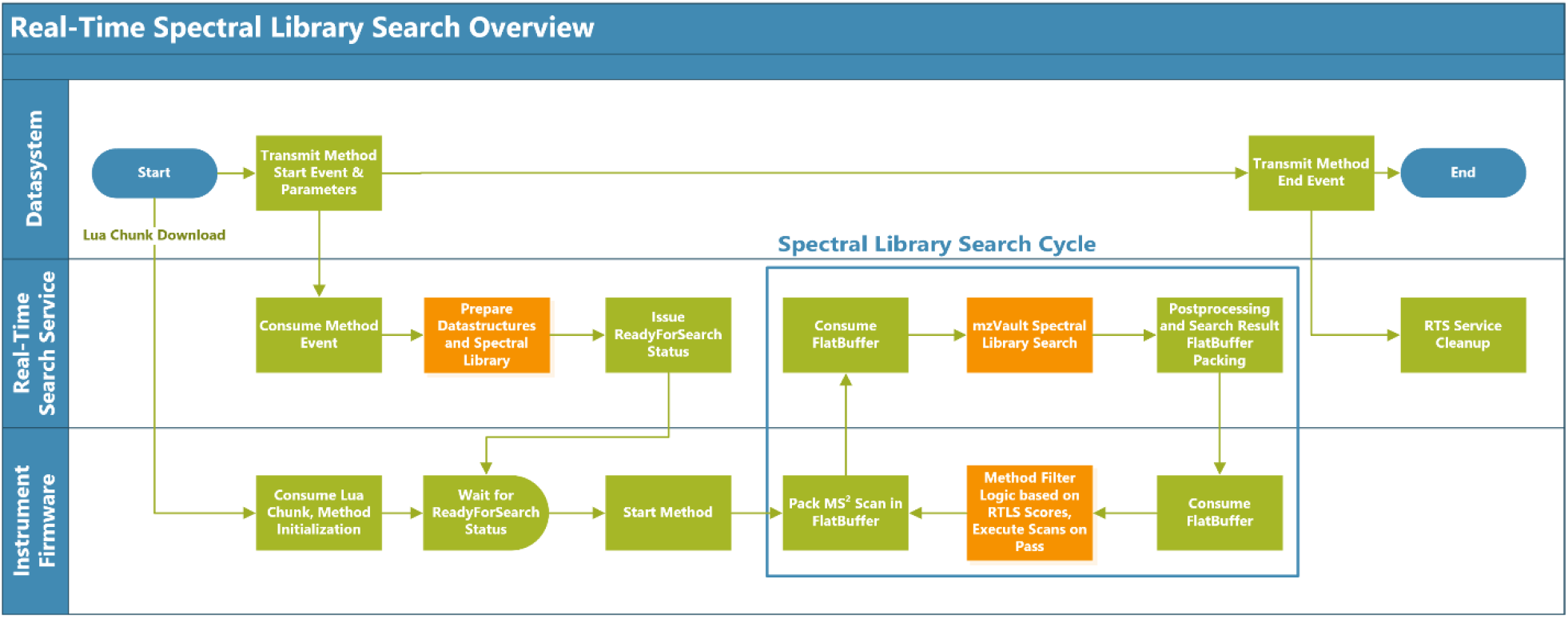
Real-Time Spectral Library Search Overview flowchart showing which systems handle method execution and the steps involved in RTLS.

Upon receipt, the Real-Time Search service queries the in-memory index for spectral candidates within the defined *m/z* tolerance boundaries relative to the assigned precursor *m/z* from the instrument. For each adduct included in the adduct search table the search service performs adduct offset queries, in which RTLS attempts to identify the analyte under the assumption that the precursor fragmented was one of the considered adducted molecular species. For each of these considered adduct forms, RTLS queries the library for spectral candidates at the calculated *m/z* of the hypothetical protonated/deprotonated form corresponding to the relevant charge state. Candidates are additionally subject to filtration based on the collision energy of the scan, and the collision energy tolerance provided within the RTLS filter parameters. If a stepped collision energy is used, the tolerance is applied to extend from the highest and lowest values of the stepped energies and all values within that range are considered as viable comparisons. Each of the spectral library entries under consideration are compared with the experimental spectrum by calculation of the cosine similarity and ranked accordingly. The highest two scoring entries considered are used to calculate their Confidence scores, as implemented within Thermo Scientific™ Mass Frontier™ software and Compound Discoverer software. The difference between the cosine and confidence scores of these two candidates are made available as the “Delta Cosine Score” and “Delta Confidence Score”, respectively. Thus, four scores can be used within each instance of the RTLS filter for selecting subsets of the scoring population: the cosine score, confidence score, and their corresponding delta scores. Each score threshold can require that values to pass be either greater than or equal to the stated value (“At Least”) or below (“Less than”) the given value. While the search is running, the instrument continues to operate normally, collecting additional MS^2^ scans in accordance with the method design. The RTLS filter additionally supports the ability to discontinue a search should the specified maximum search duration, given in milliseconds, be reached. If this limit is reached, the search aborts and the firmware receives an empty set of results for that scan.

Once the search results have been obtained, the top search result is again packed into a FlatBuffer, and its scores returned to the instrument firmware. Upon receipt, the instrument firmware will handle logic for determining if the scores constitute a set of passing values when compared against the thresholds provided by the user for each instance of the RTLS filter. Should all thresholds be met, the filter considers the search result a pass. Once a passing set of scores is obtained, the MS^2^ fragments will be considered for follow up MS^n^ scans based on use of either the “Trigger Only” parameter, in which any peak in the spectrum is considered, or if disabled, only the MS^2^ fragments which were found to match against the top scoring library candidate are considered. In newer development builds, we have enabled an option to specifically target peaks which do not match to the library candidate. Additionally, when enabled via the “Add Adducts to Dynamic Exclusion”, the calculated *m/z* values which correspond to each of the considered adduct molecular species are added to the dynamic exclusion list so that they will not be needlessly fragmented when an identification of sufficient quality has been obtained. In this design, a single scan can pass multiple RTLS filters present in a method and fail others depending on the score thresholds provided by the user. For each MS^2^ scan passed to RTLS, the Real-Time Search service will write an entry within the output tab-delimited (.tsv) file. An example of the RTLS node user interface from the method editor (Supporting Figure 1) and the results contained within the .tsv file (Supporting Table 1) can be found in the supporting information.

**Table 1.**
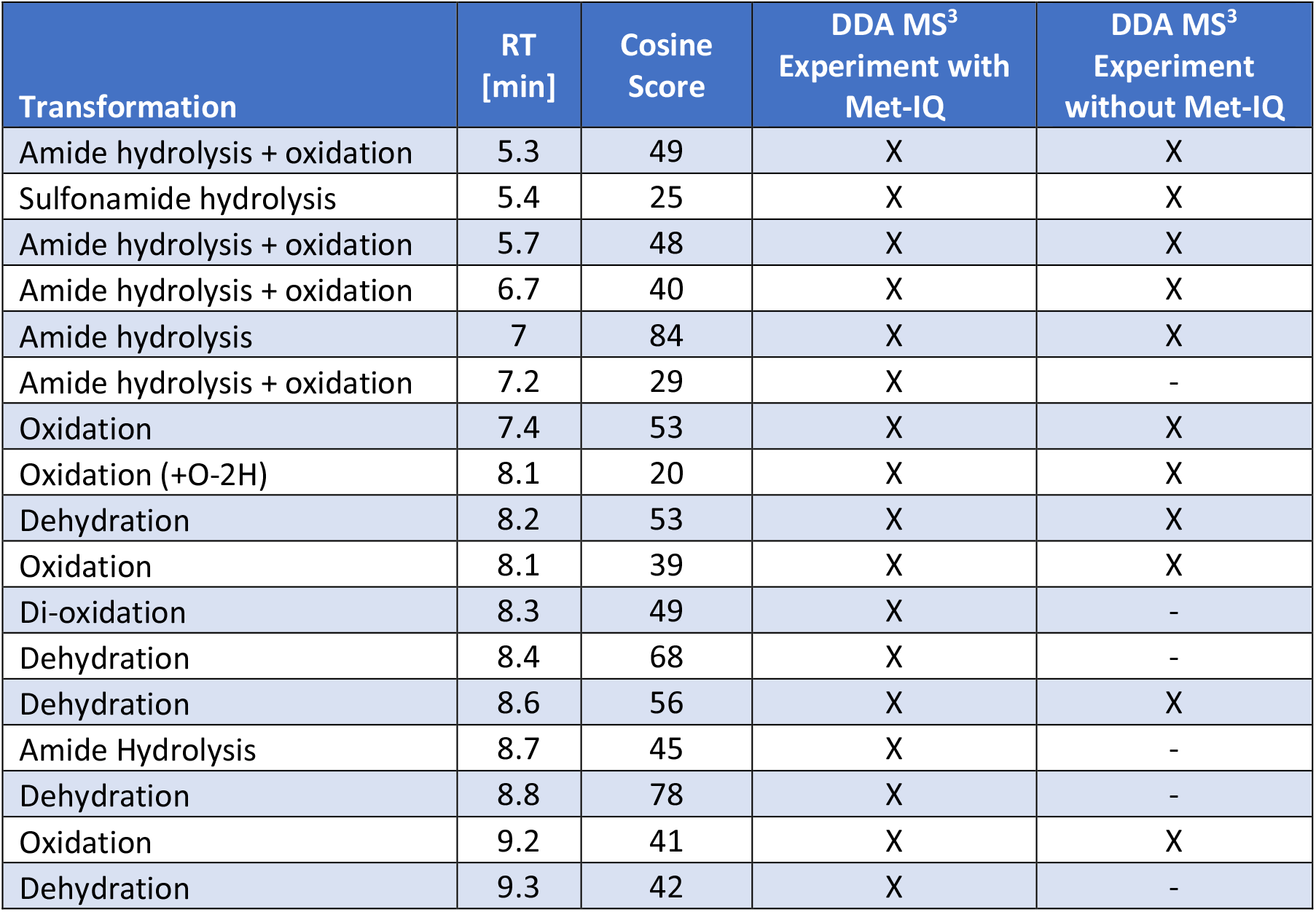
Transformation products of Amprenavir with retention time (RT), cosine score from RTLS, and whether a MS^3^ scan was triggered with or without Met-IQ.

### Impact of RTLS on Data Acquisition

The benefits of Real-Time Library Search can be broken down into two main categories. The first benefit is that the instrument precursor-selection logic is more optimized. In traditional data dependent MS^n^ acquisitions, the instrument conducts MS^n^ scans on every MS^2^ fragment that meets the triggering requirements. This leads to excess time being spent acquiring data on peaks regardless of whether they are likely compounds of interest. The Met-IQ data acquisition strategy makes use of Real-Time Library Search to restrict MS^3^ data acquisition to likely targets only.

For the digested Amprenavir sample, the instrument compared each MS^2^ spectra to the reference library containing unmetabolized Amprenavir mass spectra. When sufficient matched peaks were found, as shown in the example in Figure 2, to yield a cosine score of 15 or higher, ddMS^3^ was performed.

**Figure 2:**
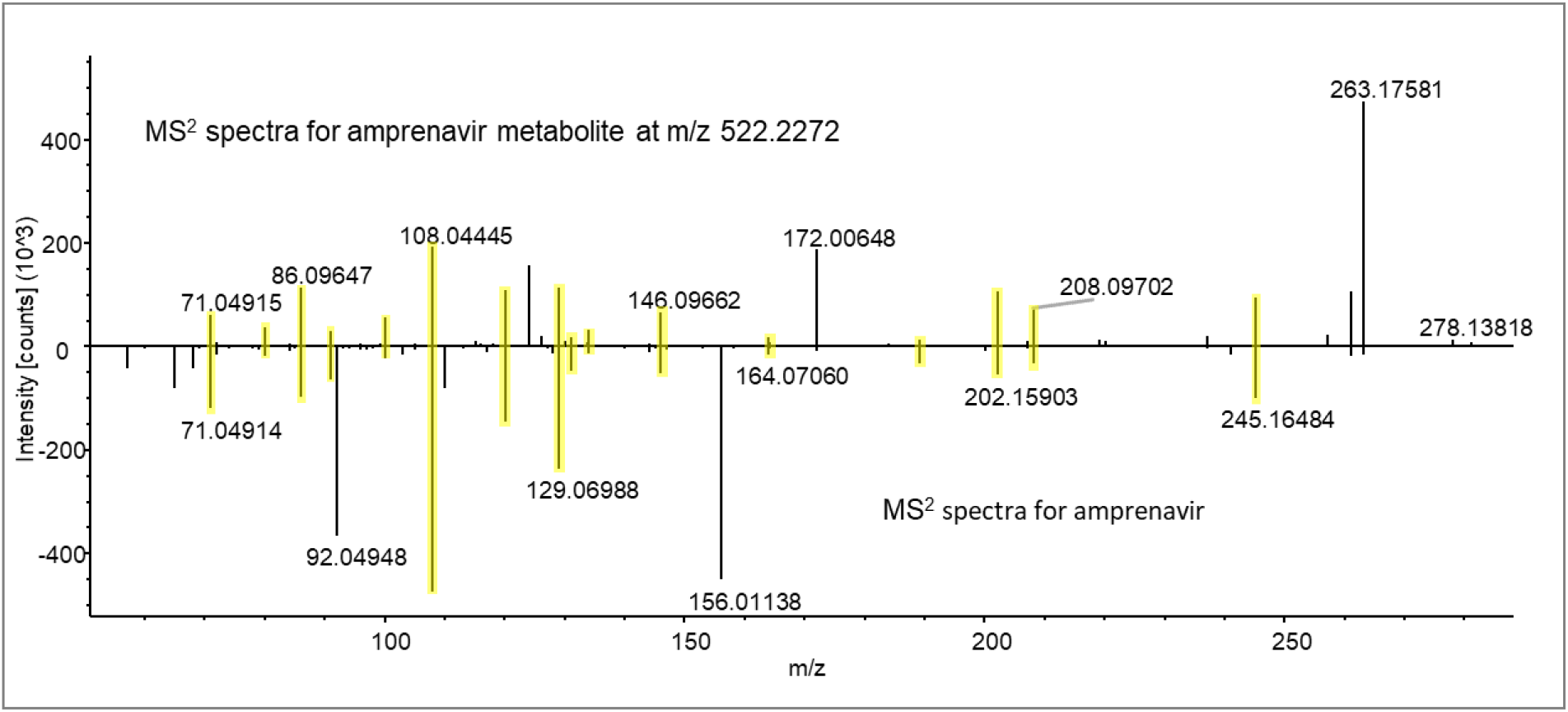
Mirror plot of MS^2^ spectra for Amprenavir metabolite and Amprenavir. Matching fragments are highlighted in yellow.

Limiting data acquisition to likely compounds of interest reduced the number of triggers for ddMS^3^ spectra which in turn enabled the instrument to spend more time sampling compounds with ddMS^2^. As shown in Figure 3, the number of MS^2^ collected in a single run increased by more than 3.4 with the use of Met-IQ relative to the DDA control, leading to an increased depth of sampling in a single run.

**Figure 3:**
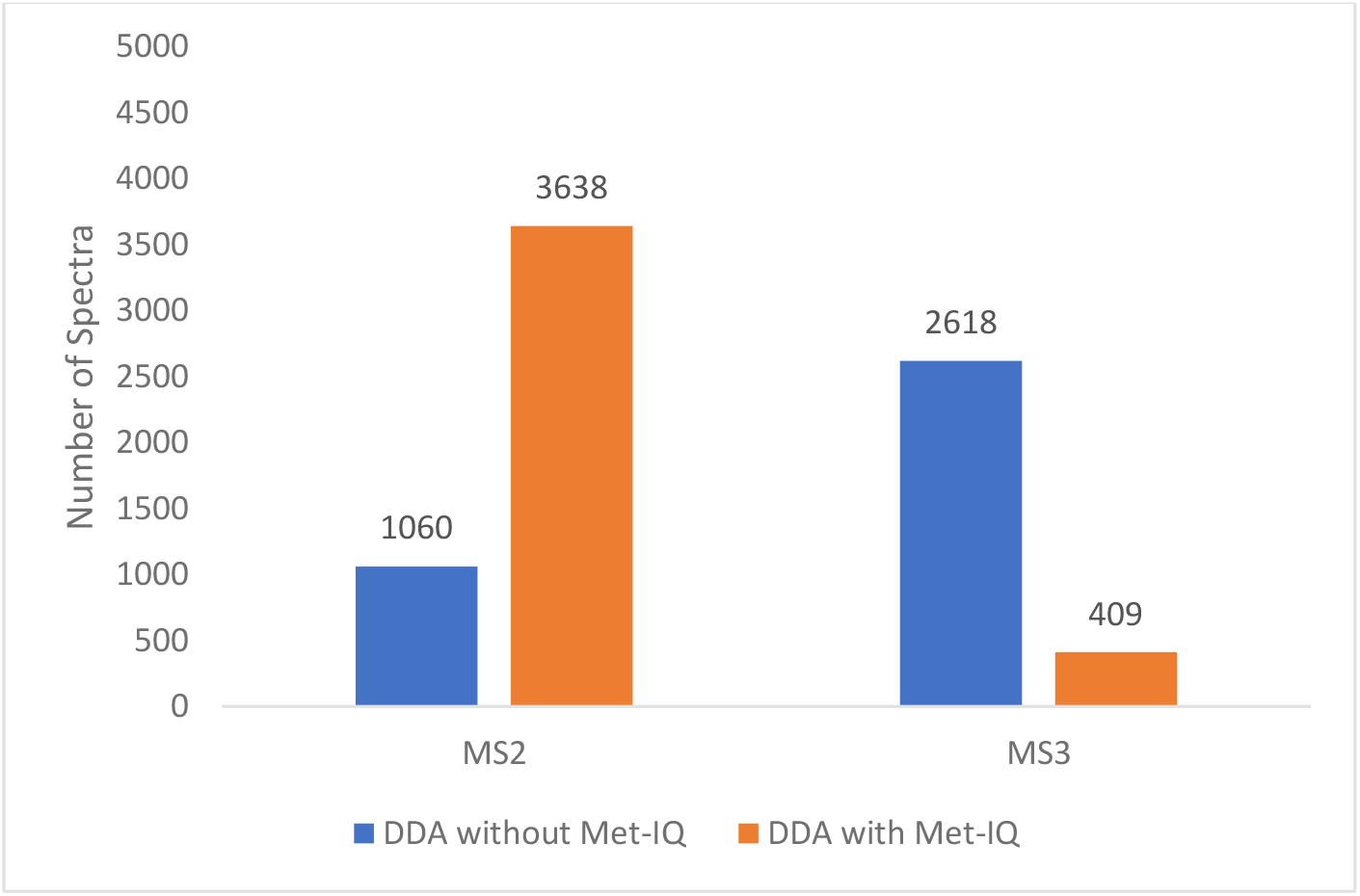
Number of MS^2^ and MS^3^ spectra collected with and without the Met-IQ. dd MS^3^ spectra were collected for the top 3 product ions with highest *m/z* for each of the RTLS-selected dd MS^2^ scans.

To evaluate the increased depth of sampling, the *m/z* values that triggered MS^3^ were compared to a list of known Amprenavir transformation products. To qualify as a potential metabolite a compound had to have the correct accurate mass of a known metabolite and have at least 1 fragment in common with the parent molecule. A total of seventeen unique metabolites were found in a single run using the Met-IQ data acquisition strategy compared to only eleven using the traditional DDA, an increase of roughly 55%, as shown in table 1.

The second benefit of the Met-IQ data acquisition strategy is the simplification of post-acquisition data analysis. In addition to increased depth of sampling there is a reduction in the raw data volume and complexity. There is over a six-fold decrease in the number of MS^3^ spectra, representing a significant decrease in sampling of unlikely targets (Fig. 3). In addition, each compound that triggered MS^3^ had at least one fragment in common with Amprenavir, flagging it as a potential metabolite. The Real-time Library Search feature is also capable of identifying MS^2^ fragments that aren’t shared with the library compound (i.e. unmatched fragments). These fragments can then be then selected for MS^3^. By focusing on unmatched fragments, this acquisition strategy increases the probability of collecting structural information about unknown portions of the molecular structure. In Figure 4A, the fragment at *m/z* 263.1758 for the Amprenavir metabolite is at a significantly higher intensity relative to the Amprenavir standard spectrum. As shown in the spectrum from Compound Discoverer software, there is no predicted structure for this fragment. However, in the MS^3^ fragmentation spectra for this fragment in Figure 4B, multiple sub-structures could be annotated by Compound Discoverer software, providing valuable structural information about the metabolite.

**Figure 4:**
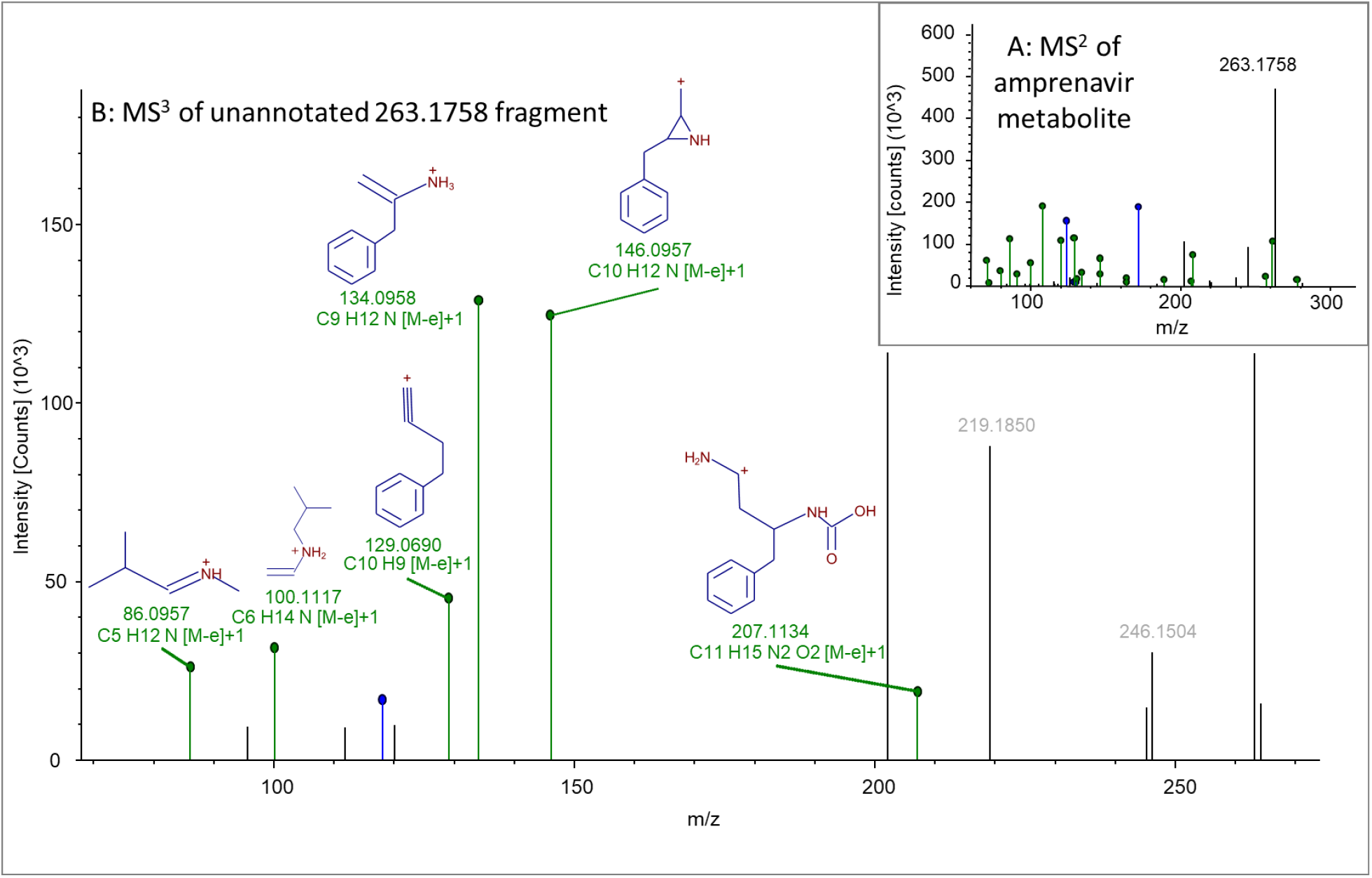
(A) The MS^2^ spectra of Amprenavir oxidation metabolite. (B) The MS^3^ spectra of the unannotated MS^2^ fragment at *m/z* 263.1758

Not shown in Table 1 is a compound corresponding to Oxidation + Cys-Gly-conjugation that was detected using traditional DDA at *m/z* 698.2524 but did not trigger MS^3^ spectra in Met-IQ experiment. The compound with the accurate mass for the oxidation and Cys-Gly-conjugation transformation product of Amprenavir didn’t share common fragments with the parent molecule at the MS^2^ level to reach a cosine score of 15 or higher and thus did not trigger MS^3^ during the Met-IQ acquisition. The Met-IQ strategy on its own would not trigger fragmentation on transformation products that have low spectral similarity with the parent molecule. However, in case of common transformations, the method can be further modified to specifically target the *m/z* of anticipated transformation product or a neutral loss associated with that transformation to be able to examine these unique cases.

## Conclusions

In the absence of thorough knowledge of potential compounds in a sample, it can be difficult to fully characterize unknowns without performing multiple analyses. In complex matrices, compounds can undergo unique and unexpected transformations and lead to samples that are difficult to analyze. In depth characterization using multiple analyses yields complicated data sets that require extensive time and expertise to process. We have presented a new data acquisition strategy called Met-IQ in which Real-Time Library Search is used to direct data acquisition towards likely analytes of interest to yield data sets with more depth of sampling and less complexity. Data dependent MS^3^ analysis was carried out on a metabolized sample of the drug Amprenavir where putative metabolites were targeted based on their spectral similarity to the parent drug. Experimental MS^2^ were compared to library spectra in real time and compounds exhibiting shared fragments were further analyzed. By limiting MS^3^ analysis only to the targets that displayed similarity with the parent drug, the number of MS^2^ spectra collected increased 3.4-fold and the number of metabolites found in a single run increased by 55% relative to conventional data dependent analysis. Additional structural information was also acquired by automatically targeting MS^2^ fragments of experimental compounds that weren’t found in the library spectra. Out of the 18 potential transformation products found in these experiments, only a single product was missed using the Met-IQ data acquisition strategy due to a lack of spectral similarity to the parent drug. However, this type of transformation can be found by making use of alternate scan filters available in the method editor software. This approach to identifying unknown small molecule species based on their spectral similarity to the known structure can be further applied to screen for unknown compounds belonging to a specific compound class in many analytical fields, including analysis extractables and leachables, clinical toxicology, environmental and food safety and others.

## Acknowledgment

We would like to thank Dr. Shunguang Ma from Genentech (South San Francisco, CA) for providing metabolized Amprenavir sample; Mark Sanders, Jim Yano, Romain Huguet, Thomas Moehring and Tim Stratton from ThermoFisher Scientific for helpful discussions.

For General Lab Use Only. Not for use in diagnostic procedures.

