## Supplemental Information for "Novel Real-Time Library Search driven data acquisition strategy for identification and characterization of metabolites"

Detailed RTLS Parameters for Met-IQ

Collision energy tolerance was set so only library spectra collected with collision energy within ±15 of the experimental spectra would be considered. To ensure all experimental MS^2^ spectra were compared against the amprenavir spectra to check for similar MS^2^ fragments, the precursor search tolerance was set arbitrarily large at 50,000,000 ppm. Under the adduct masses table the adduct molecular species and charge state were left at the default values of “M” and “0” which considers all protonated forms of the library compound at any charge state. The maximum search time was left at the default value of 150 milliseconds. For non-matched peaks (peaks found experimentally but not in the spectral library) to be considered, “Use as Trigger Only” was set to on. All adducts were considered during this experiment so “Add Adducts to Dynamic Exclusion” was set to off.

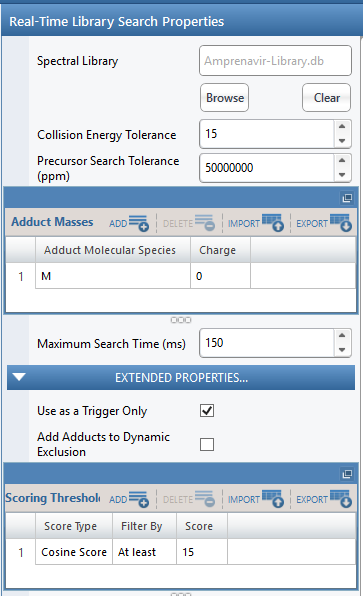

Supporting Figure 1: User interface of Real-Time Library Search filter

Contents of the .tsv File

Any time a method containing the RTLS filter is run, a .tsv text file is generated with the results of the Real-Time Library Search. A segment from a .tsv file that was copied over to an excel document is found in Supporting Table 1. For each MS^2^ scan evaluated, the .tsv file records the scan number, isolation m/z, isolation z (not shown below as it was always 1 for this experiment), formula for the library match (not shown below because the library only contained amprenavir), compound class of the library match (not used in this experiment), charge state (not shown below as it was always 1 for this experiment), compound Name from the library match, search time in milliseconds, cosine score, confidence score, cosine delta score, confidence delta score, the delta ppm in mass between the experimental m/z and library match, and the candidate count (the number of candidates from the library that were considered valid comparisons based on the search criteria in the RTLS node). If the cosine score, confidence score, cosine delta score, and confidence delta scores meet the criteria set (if that particular score has a criteria) in the RTLS node, then the node will pass that scan along to trigger a scan event on that compound.

Supporting Table 1: Example information that is captured in the .tsv file generated when a method containing RTLS is run.

| Scan Number | Isolation m/z | Compound Name | Search Time (ms) | Cosine Score | Confidence Score | Cosine Delta Score | Confidence Delta Score |
| --- | --- | --- | --- | --- | --- | --- | --- |
| 155 | 348.0135 | Amprenavir | 7 | 11 | 34 | 11 | 34 |
| 159 | 744.0829 | Amprenavir | 4 | 29 | 17 | 29 | 17 |
| 161 | 335.0193 | Amprenavir | 8 | 15 | 44 | 15 | 44 |
| 164 | 744.0829 | Amprenavir | 6 | 29 | 17 | 29 | 17 |
| 165 | 346.0094 | Amprenavir | 5 | 10 | 43 | 10 | 43 |
